# The face module emerged in a deep convolutional neural network selectively deprived of face experience

**DOI:** 10.1101/2020.07.06.189407

**Authors:** Shan Xu, Yiyuan Zhang, Zonglei Zhen, Jia Liu

## Abstract

Can we recognize faces with zero experience on faces? This question is critical because it examines the role of experiences in the formation of domain-specific modules in the brain. Investigation with humans and non-human animals on this issue cannot easily dissociate the effect of the visual experience from that of the hardwired domain-specificity. Therefore the present study built a model of selective deprivation of the experience on faces with a representative deep convolutional neural network, AlexNet, by removing all images containing faces from its training stimuli. This model did not show significant deficits in face categorization and discrimination, and face-selective modules automatically emerged. However, the deprivation reduced the domain-specificity of the face module. In sum, our study provides undisputable evidence on the role of nature versus nurture in developing the domain-specific modules that domain-specificity may evolve from non-specific experience without genetic predisposition, and is further fine-tuned by domain-specific experience.

## 1 Introduction

A fundamental question in cognitive neuroscience is how nature and nurture form our cognitive modules. In the center of the debate is the origin of face recognition ability. Numerous studies have revealed both behavioral and neural signatures of face-specific processing, indicating a face module in the brain (for reviews, see Freiwald, Duchaine, & Yovel, 2016; Kanwisher & Yovel, 2006). Further studies from behavioral genetics revealed the contribution of genetics on the development of the face-specific recognition ability in humans (Wilmer et al., 2010; Zhu et al., 2010). Collectively, these studies suggest an innate domain-specific module for face cognition. However, it is unclear whether the visual experience is also necessary for the development of the face module.

A direct approach to address this question is visual deprivation. Two studies on monkeys selectively deprived the visual experience of faces since birth, while leaving the rest of experiences untouched (Arcaro, Schade, Vincent, Ponce, & Livingstone, 2017; Sugita, 2008). They report that face-deprived monkeys are still capable of categorizing and discriminating faces (Sugita, 2008), though less prominent in selective looking preference to faces over non-face objects (Arcaro et al., 2017). Further examination of the brain of the experience-deprived monkeys fails to localize typical face-selective cortical regions with the standard criterion; however, in the inferior temporal cortex where face-selective regions are normally localized, face-selective activation (i.e., neural responses to faces larger than nonface objects) is observed (Arcaro et al., 2017). Taken together, without visual experiences of faces, rudimental functions to process faces may still evolve to some extent.

Two related but independent hypotheses may explain the emergence of the face module without face experiences. An intuitive answer is that the rudimental functions are hardwired in the brain by genetic predisposition (McKone, Crookes, Jeffery, & Dilks, 2012; Wilmer et al., 2010). Alternatively, we argue that the face module may emerge from experiences on nonface objects and related general-purpose processes, because representations for faces may be constructed by abundant features derived from nonface objects. Unfortunately, studies on humans and monkeys are unable to thoroughly decouple the effect of nature and nurture to test these two hypotheses.

Recent advances in deep convolutional neural network (DCNN) provide an ideal test platform to examine the impact of visual experiences on face modules without genetic predisposition. DCNNs are found similar to human visual cortex both structurally and functionally (Kriegeskorte, 2015), but free of any predisposition on functional modules. Therefore, with DCNNs we can manipulate experiences without considering interactions from genetic predisposition. In this study, we asked whether DCNNs can achieve face-specific recognition ability when visual experiences on faces were selectively deprived.

To do this, we trained a representative DCNN, AlexNet (Krizhevsky, Sutskever, & Hinton, 2012), to categorize nonface objects with face images carefully removed from the training dataset. Once this face-deprived DCNN (d-AlexNet) was trained, we compared its behavioral performance to that of a normal AlexNet of the same architecture but with faces present during training in both face categorization (i.e., differentiating faces from nonface objects) and discrimination (i.e., discriminating faces among different individuals) tasks. We predicted that the d-AlexNet, though without predisposition and experiences of faces, may still develop face selectivity through its visual experiences of nonface objects.

## 2 Results

The d-AlexNet was trained with a dataset of 662,619 non-face images consisting of 736 non-face categories, generated by removing images containing faces from the ILSVRC 2012 dataset (Figure 1a). The d-AlexNet was initialized and trained in the same way as the AlexNet, and the resultant top-1 accuracy (57.29%) and the top-5 accuracy (80.11%) were comparable with the pre-trained AlexNet.

**Figure 1.**
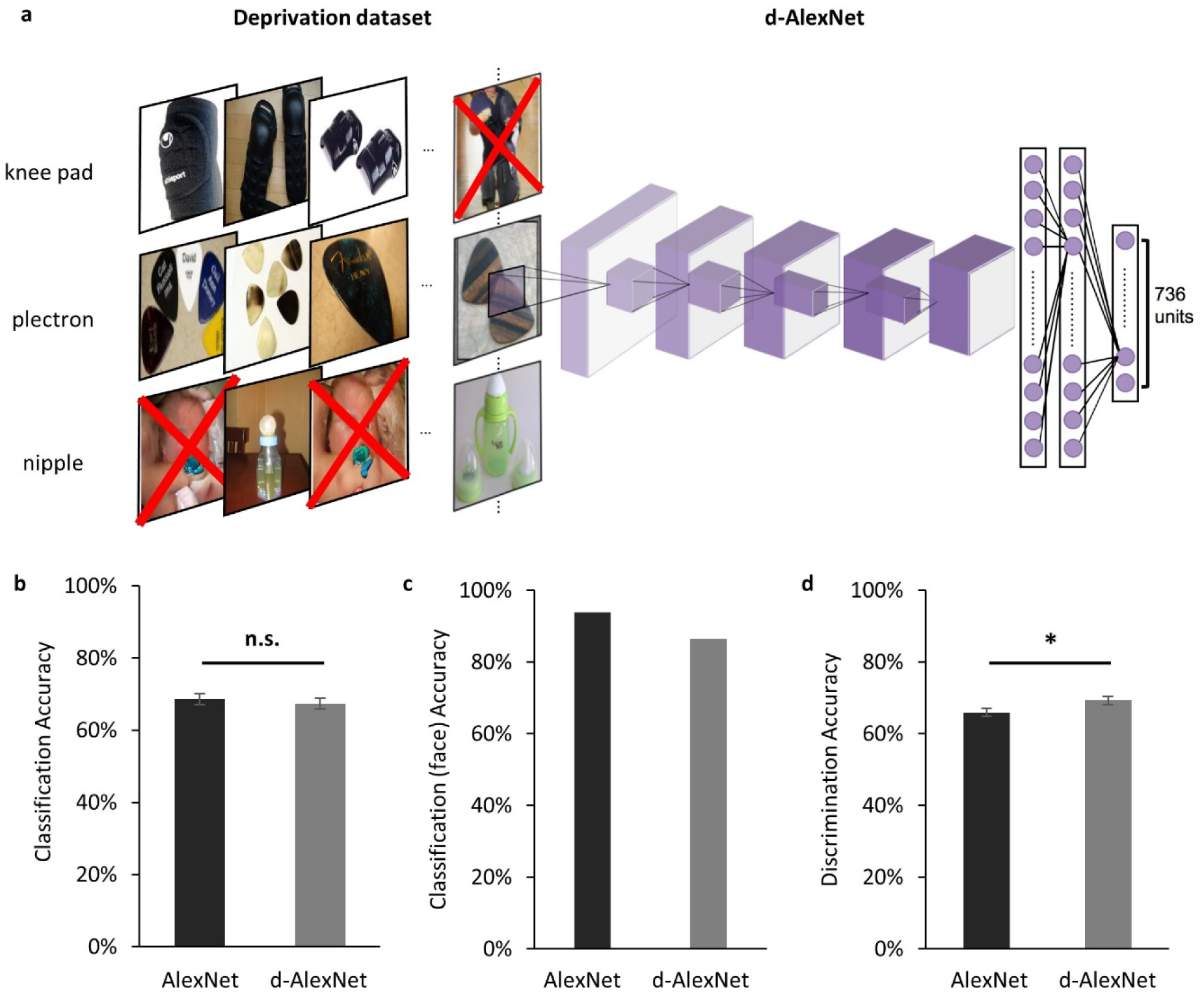
(a) An illustration of the screening to remove images containing faces for the d-AlexNet. The ‘faces’ shown in the figure were AI-generated for illustration purpose only, and therefore have no relation to real person. In the experiment, face images were from the ImageNet, with real persons’ faces. (b) The classification performance across categories of the two DCNNs was comparable. (c) Both DCNNs achieved high accuracy in categorizing faces from other images. (d) Both DCNNs’ performance in discriminating faces was above the chance level, and the d-AlexNet’s accuracy was significantly higher than that of the AlexNet. The error bar denotes standard error. The asterisk denotes statistical significance (α = .05). n.s. denotes no significance.

We first examined the performance of the d-AlexNet in two representative tasks of face processing, face categorization (i.e., differentiating faces from non-face objects) and face discrimination (i.e., identifying different individuals). The output of Conv5 after ReLU of the d-AlexNet was used to classify objects in the classification dataset (see Materials and Methods). The averaged categorization accuracy of the d-AlexNet (67.40%) was well above the chance level (0.49%), and comparable to that in the AlexNet (68.60%, *t* (204) = 1.26, *p* = 0.209, Cohen’s d = 0.007, Figure 1b). Critically, the d-AlexNet, although with no experience on faces, succeeded in the face categorization task, with an accuracy of 86.50% in categorizing faces from non-face objects. Note that the accuracy was numerically smaller than the AlexNet’s accuracy in categorizing faces (93.90%) though (Figure 1c).

A similar pattern was observed in the face discrimination task. In this task, the output of Conv5 after ReLU of each DCNN was used to identify 33,250 face images into 133 identities in the discrimination dataset (see Materials and Methods). As expected, the AlexNet was capable of face discrimination (65.9%), well above the chance level (0.75%), consistent with previous studies (AbdAlmageed et al., 2016; Grundstrom, Chen, Ljungqvist, & Astrom, 2016). Critically, the d-AlexNet also showed the capability of discriminating faces, with an accuracy of 69.30% that was even significantly higher than that of the AlexNet, *t* (132) = 3.16, *p* = .002, Cohen’s d = 0.20, (Figure 1d). Taken together, visual experiences on faces seemed not necessary for developing basic functions of processing faces.

Was a face module formed in the d-AlexNet to support these functions? To answer this question, we searched all the channels in Conv5 of the d-AlexNet, where face-selective channels have been previously identified in the AlexNet (Baek, Song, Jang, Kim, & Paik, 2019). To do this, we calculated the activation of each channel in Conv5 after ReLU in response to each category of the classification dataset, and then identified channels that showed significantly higher response to faces than non-face images with Mann-Whitney U test (*p*s < .05, Bonferroni corrected). Two face-selective channels (Ch29 and Ch50) met this criterion in the d-AlexNet (for an example channel, see Figure 2a, right), whereas four face-selective channels (Ch195, Ch125, Ch60, and Ch187) were identified in the AlexNet (for an example channel, see Figure 2a, left). The face-selective channels in two DCNNs differed in selectivity. The averaged selective ratio, the ratio of the activation magnitude to faces by that to the most activated non-face object category, was 1.29 (range: 1.22 - 1.36) in the d-AlexNet, much lower than that in the AlexNet (average ratio: 3.63, range: 1.43 - 6.66). The lifetime sparseness, which measures the breadth of tuning of a channel in response to a set of categories, also showed a similar result. The average lifetime sparseness index of the face channels in the AlexNet (mean = 0.25, range: 0.11 – 0.51) was smaller than that in the d-AlexNet (mean = 0.71, range: 0.70 – 0.71), indicating higher face selectivity in the AlexNet than that in the d-AlexNet. Taken together, this finding suggested that the face-selective channels already emerged in the d-AlexNet, though the face selectivity was weaker.

**Figure 2.**
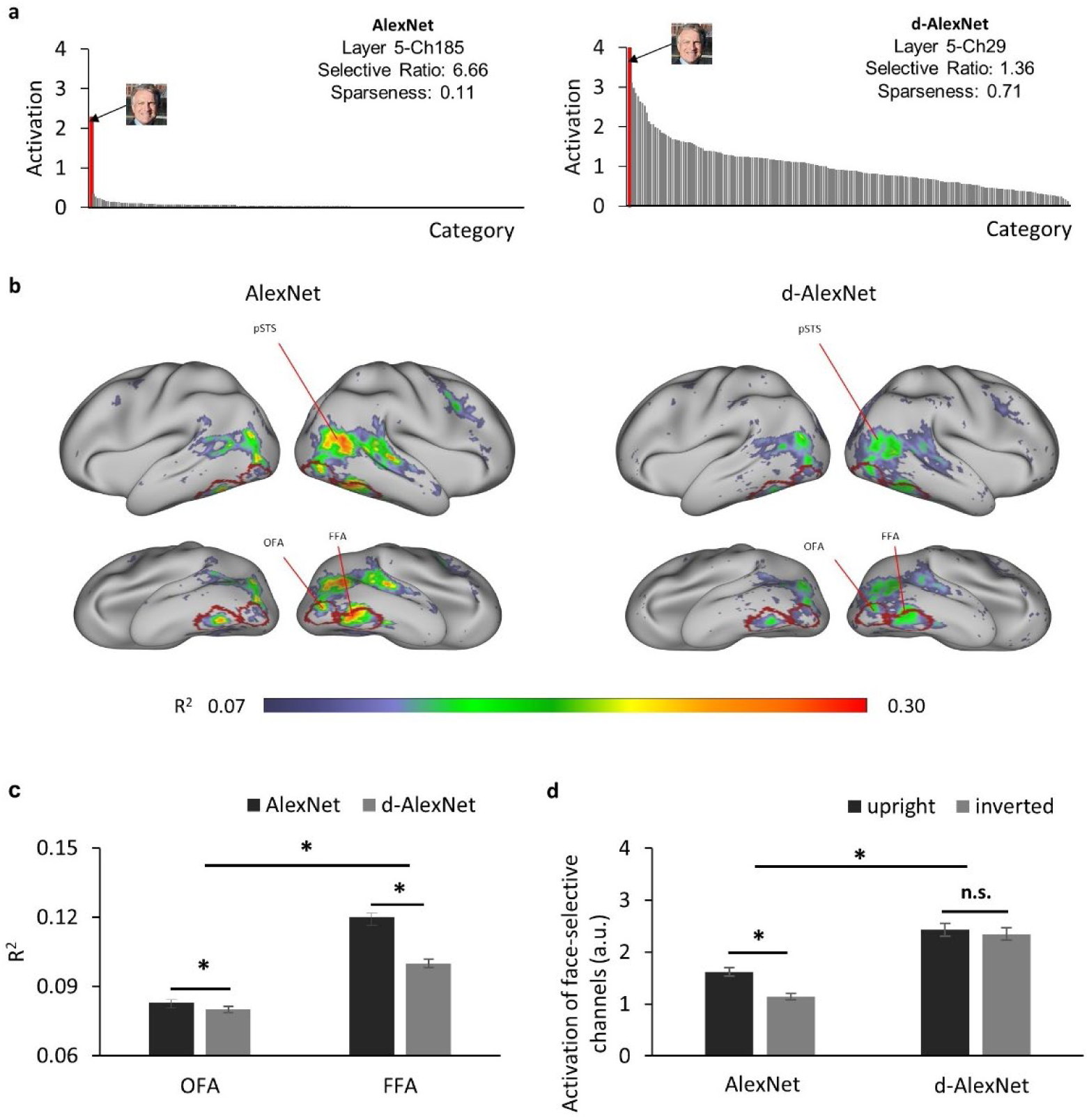
(a) The category-wise activation profiles of example face-selective channels of the AlexNet (left) and the d-AlexNet (right). The ‘faces’ shown here were AI-generated for illustration purpose only. (b) The R^2^ maps of the regression with the activation of the d-AlexNet’s (right) or the AlexNet’s face-selective channels (left) as the independent variables. The higher R^2^ in multiple regression, the better correspondence between the face channels in the DCNNs and the face-selective regions in the human brain. The crimson lines delineate the ROIs of the OFA and the FFA. (c) The face-channels of both DCNNs corresponded better with the FFA than the OFA, and the difference between the AlexNet and the d-AlexNet was larger in the FFA. (d) Face inversion effect. The average activation amplitude of the top two face-selective channels differed in response to upright and inverted faces in the AlexNet but not the d-AlexNet. The error bar denotes standard error. The asterisk denotes statistical significance (α = .05). n.s. denotes no significance.

How did the face-selective channels correspond to face-selective cortical regions in humans, such as the FFA and OFA? To answer this question, we calculated the coefficient of determination (R^2^) of the multiple regression with the output of the face-selective channels as regressors and the fMRI signals from human visual cortex in response to movies on natural vision as the regressand (see Materials and Methods). As shown in Figure 2b (right), the face-selective channels identified in the d-AlexNet corresponded to the bilateral FFA, OFA, and the posterior superior temporal sulcus face area (pSTS-FA). Similar correspondence was also found with the top two face-selective channels in the AlexNet (Figure 2b, left). Direct visual inspection revealed that the deprivation weakened the correspondence between the face-selective channels and face-selective regions in human brain. This observation was confirmed by the main effect of visual experiences (*F* (1,798) = 161.97, *p* < .001, partial η^2^ = 0.17) in a two-way ANOVA of visual experiences (d-AlexNet vs. AlexNet) by regional correspondence (the OFA versus the FFA). In addition, the main effect of the regional correspondence showed that the response profile of the face-selective channels in the DCNNs fitted better with the activation of the FFA than that of the OFA (*F* (1,798) = 98.69, *p* = .001, partial η^2^ = 0.11), suggesting that the face-selective channels in DCNNs may in general prefer to process faces as a whole than face parts. Critically, the two-way interaction was significant (*F* (1,798) = 84.9, *p* < .001, partial η^2^ = 0.10), indicating that the experience affected the correspondence to the FFA and OFA disproportionally. A simple effect analysis revealed that the correspondence to the FFA (MD = 0.023, *p* < .001) was increased by face-specific experiences to a significantly larger extent than that to the OFA (MD = 0.004, *p* = .013, Figure 2c). Since the FFA is more involved in holistic processing of faces and the OFA is more dedicated to the part-based analysis, the disproportional decrease in correspondence between the face-selective channels in the d-AlexNet and the FFA implied that the role of the experience on faces was to facilitate the processing of faces as a whole.

To test this conjecture, we examined how the d-AlexNet responded to inverted faces, a behavioral signature of face-specific processing. As expected, there was a face inversion effect in the AlexNet’s face-selective channels, with the magnitude of the activation to upright faces significantly larger than that to inverted faces (*t* (19) = 6.45, *p* < .001, Cohen’s d = 1.44) (Figure 2d). However, no inversion effect was observed in the d-AlexNet, as the magnitude of the activation to upright faces was not significantly larger than that to inverted faces (*t* (19) = 0.86, *p* = .40). The lack of the inversion effect in the d-AlexNet was further supported by a two-way interaction of visual experience by orientation of faces, *F* (1, 19) = 7.79, *p* = .012, partial η^2^ = 0.29. That is, unlike the AlexNet, the d-AlexNet processed upright faces in the same fashion as inverted faces.

Previous studies on human suggested that inverted faces are processed in an object-like fashion. That is, it relies more on the parts-based analysis than the holistic processing. Therefore, we speculated that in the d-AlexNet faces were also represented more like non-face objects. To test this speculation, we first compared the representational similarity among responses in Conv5 to faces and bowling-pins, a novel object category that was not exposed to either DCNNs. As expected, the two-way interaction of experience (AlexNet versus d-AlexNet) by category (faces versus bowling-pins) was significant (*F* (1, 6,318) = 4,110.88, *p* <.001, partial η^2^ = 0.39), and the simple effect analysis suggested that the representation for faces in the AlexNet was more similar between each other than in the d-AlexNet (MD = 0.16, *p* <.001), whereas the within-category representation similarity for bowling-pins showed the same but numerically smaller between-DCNN difference (MD = 0.005, *p* = .002) (Figure 3a).

**Figure 3.**
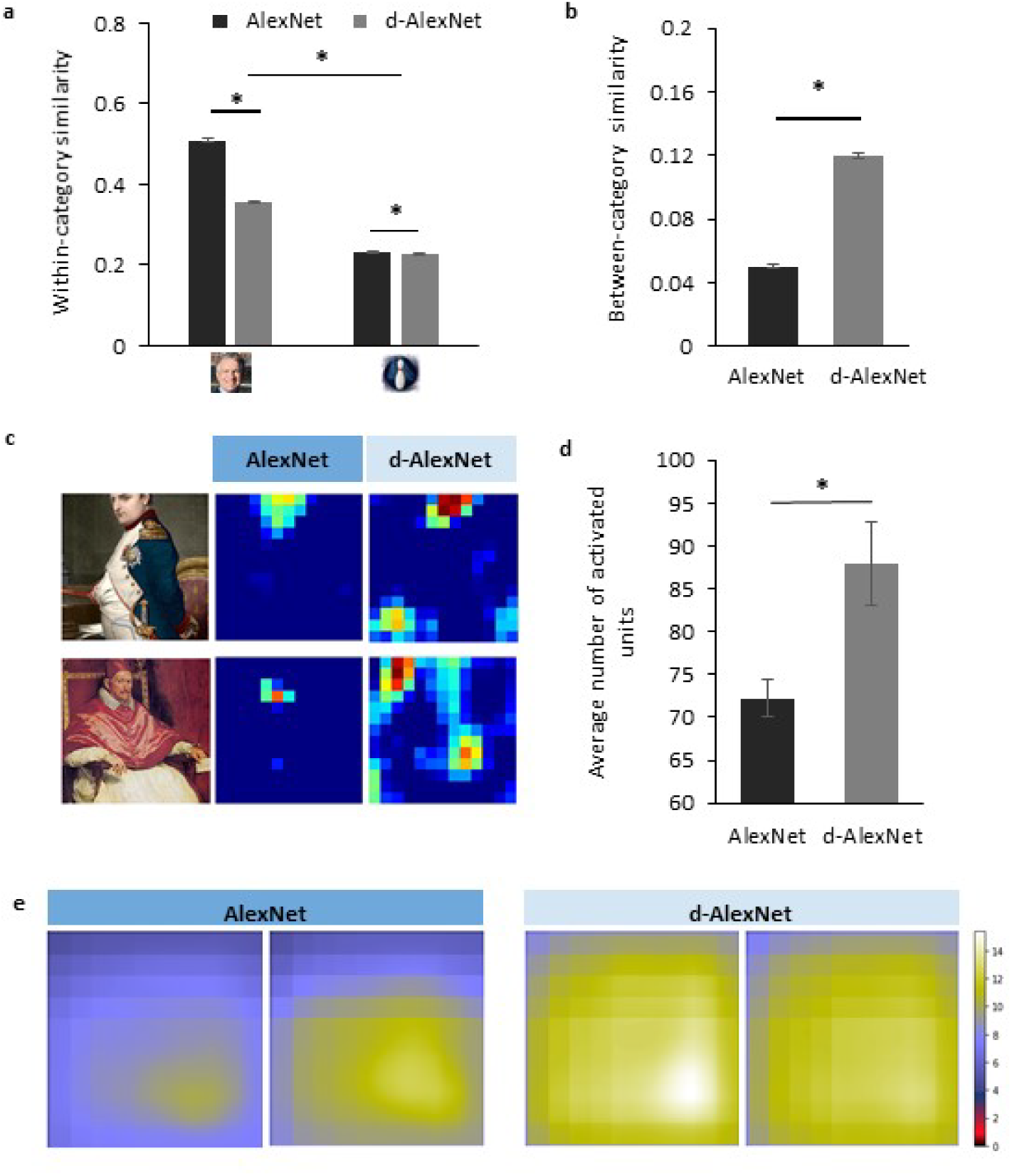
(a) The within-category similarity in the face category and an unseen non-face category (bowling pins) in the DCNNs. (b) The between-category similarity between faces and bowling pins. (c) The activation maps of a typical face-selective channel of each DCNN in responses to natural images containing faces. Each pixel denotes activation in one unit. The images shown here were historical portrait paintings downloaded from the Internet for illustration purpose only, and are different from the images used in this study. (d) The extent of activation of the face-selective channels of each DCNN in responses to natural images containing faces. (e) The empirical receptive fields of the face-selective channels of each DCNN. The error bar denotes standard error. The asterisk denotes statistical significance (α = .05).

A more critical test was to examine how face-specific experiences made faces being processed differently from objects. Here we calculated between-category similarities between faces and bowling-pins. We found that the between-category similarity between faces and bowling-pins was significantly higher in the d-AlexNet than that in the AlexNet (*t* (3,159) = 42.42, MD = 0.07, *p* < .001, Cohen’s d = 0.76) (Figure 3b), suggesting that faces in the d-AlexNet were indeed represented more like objects. In short, although d-AlexNet was able to perform face tasks similar to the one with face-specific experiences, it represented faces in an object-like fashion.

Finally, we asked how faceness was achieved in DCNNs with face-specific experiences. Neurophysiological studies on monkeys demonstrate experience-associated sharpening of neural response, with fewer neurons activated after learning. Here we performed a similar analysis by measuring the number of non-zero units (i.e., units with above-zero activation) of the face-selective channels activated by natural images containing faces. As shown in the activation map (Figure 3c), a smaller number of units were activated by faces in the AlexNet than that in the d-AlexNet (*t* (19) = 3.317, MD =15.78, Cohen’s d = 0.74) (Figure 3d), suggesting that the experience on faces made the representation to faces sparser, and thus more effective. Another effect of visual experiences observed in neurophysiological studies is that experiences reduce the size of neurons’ receptive field. Here we also mapped the empirical receptive field of the face-selective channels (see Materials and Methods). Similarly, we found that the empirical receptive field of the AlexNet was smaller than that of the d-AlexNet. That is, within the theoretical receptive field, the empirical receptive field of the face-selective channels in the AlexNet was tuned to focus on a smaller region by face-specific experiences (Figure 3e).

## 3 Discussion

This study presented a DCNN model of selective visual deprivation of faces. We found that without genetic predisposition and face-specific visual experiences, DCNNs were still capable of face perception. In addition, face-selective channels were also present in the d-AlexNet, which corresponded to human face-selective regions. That is, the visual experience of faces was not necessary for an intelligent system to develop a face-selective module. On the other hand, besides the slightly compromised selectivity of the module, the deprivation led the d-AlexNet to process faces in a fashion more similar to that of processing objects. Indeed, face-inversion effect was absent in the d-AlexNet, and the representation of faces was more similar to objects as compared to the AlexNet. Finally, face-specific experiences might affect face processing by fine-tuning the sparse coding and the size of the receptive field of the face-selective channels. In sum, our study addressed a long-standing debate on nature versus nurture in developing the face-specific module, and illuminated the role of visual experiences in shaping the module.

It may seems surprising that without domain-specific visual experience, the face-selective processing and module still emerged in the d-AlexNet; yet this finding is consistent with previous studies on non-human primates and new-born human infants (Bushneil, Sai, & Mullin, 1989; Sugita, 2008; Valenza, Simion, Cassia, & Umiltà, 1996), where the face-specific experience is found not necessary for face detection and recognition. However, the experience-independent face processing is largely attributed to either innate face-specific mechanisms (McKone et al., 2012; Morton & Johnson, 1991) or domain-general processing with predisposed biases (Simion & Di Giorgio, 2015; Simion, Macchi Cassia, Turati, & Valenza, 2001). Our study argued against this conjecture, because unlike any biological system, DCNNs have no domain-specific genetic inheritance or processing biases. Therefore, the face-specific processing observed in DCNNs had to derive from domain-specific factors.

We speculated that the face module in the d-AlexNet may result from the rich features represented in the multiple layers of the network, with which face-like features were selected to construct face-specific module. In fact, previous studies on DCNNs have shown that DCNN’s lower layers showed sensitivity to myriad visual features similar to primates’ primary visual cortex (Krizhevsky et al., 2012), while the higher layers are tuned to complex features resembling those represented in the ventral visual pathway (Güçlü & van Gerven, 2015; Yamins et al., 2014). With such repertoire of rich features, a representational space for faces, or for any natural object, may be constructed by selecting features that are potentially useful in face tasks.

Supporting evidence for this conjecture came from the observation that the d-AlexNet processed faces in an object-like fashion. For example, the face inversion effect, a behavioral signature of face-specific processing in human (Kanwisher, Tong, & Nakayama, 1998; Yin, 1969) was absent in the d-AlexNet. That is, similar to inverted faces, upright faces may also be processed like objects in the d-AlexNet. A more direct illustration of the object-like representation of faces came from the analysis on the representational similarity between faces and objects. As compared to the AlexNet, faces in the representational space of the d-AlexNet were less congregated among each other; instead they were more intermingled with non-face object categories. The finding that face representation was no longer qualitatively different from object representation may help explaining the performance of the d-AlexNet. Because faces were less segregated from objects in the representational space, the d-AlexNet’s accuracy of face categorization was worse than that of the AlexNet. In contrast, within the face category, individual faces were less congregated in the representational space; therefore, the discrimination of individual faces became easier instead, suggested by the slightly higher face discrimination accuracy in the d-AlexNet than the AlexNet. In short, when the representational space of the d-AlexNet was formed exclusively based on features from non-face stimuli, faces were represented no longer qualitatively different from non-face objects, which inevitably led to ‘object-like’ face processing.

The face-specific processing is likely achieved through prior exposure to faces. At first glance, the effect of face-specific experiences seemed quantitative, as in the AlexNet, both the selectivity to faces and the number of the face-selective channels were increased, and the correspondence between the face-selective channels and the face-selective regions in human brain was tighter. However, careful scrutiny of the difference between the two DCNNs revealed that the changes led by the experience may be qualitative. For example, the deprivation of visual experiences disproportionally weakened the DCNN-brain correspondence in the FFA as comparing to the OFA, and the FFA is engaged more in the configural processing and the OFA in parts-based analysis (Liu, Harris, & Kanwisher, 2010; Nichols, Betts, & Wilson, 2010; Zhao et al., 2014). Therefore, the ‘face-like’ face processing may come from the fact that face-specific experiences led the representation of faces more congregated within face category and more separable from the representation of non-face objects stimuli (see also Gomez, Barnett, & Grill-Spector, 2019). In this way, a relative encapsulated representation may help developing a unique way of processing faces, qualitatively different from non-face objects.

The advantage of the computational transparency of DCNNs may shed light on the development of domain specificity of the face module. First, we found that face-specific experiences increased the sparseness of face representation, as fewer units of the face channels were activated by faces in the AlexNet. The experience-dependent sparse coding has been widely discovered in the visual cortex (for reviews, see Desimone, 1996; Grill-Spector, Henson, & Martin, 2006). The experience-induced increase of sparseness is thought to reflect a preference-narrowing process that tunes neurons to a smaller range of stimuli (Kohn & Movshon, 2004); therefore, with sparse coding faces are less likely to be intermingled with non-face objects, which may lead to more congregated representations in the representational space in the AlexNet, as compared to the d-AlexNet. Second, we found that the empirical receptive field of the face channel in the AlexNet was smaller than that in the d-AlexNet, suggesting that the visual experience on faces decreased the size of the receptive field of the face channels. This finding fits perfectly with neurophysiological studies that the size of receptive fields of visual neurons is reduced after eye-opening (Braastad & Heggelund, 1985; Cantrell, Cang, Troy, & Liu, 2010; Tavazoie & Reid, 2000). Importantly, along with the refined receptive fields, the selectivity of neurons increases (Spilmann, 2014), possibly because neurons can avoid distracting information by focusing on a more restricted part of stimuli, which may further allowed finer representation of the selected regions. This is especially important for processing faces because faces are highly homogeneous, and some information is identical across faces, such as parts composition (eyes, noses, and mouth) and their configural arrangements. Therefore, the reduced receptive field of the face channels may facilitate selective analyses of discriminative face features while avoiding irrelevant information. Further, the sharpening of the receptive field and the fine-tuned selectivity may result in superior discrimination ability on faces, and allow faces to be processed at the sub-ordinate level (i.e., identification), whereas the rest of objects are largely processed at the basic level (i.e., categorization).

It has long been assumed that domain-specific visual experiences and inheritance are the pre-requisites in the development of the face module. In our study with DCNN as a model, we completely decoupled the genetic predisposition and face-specific visual experiences, and found that the representation for faces can be constructed with features from non-face objects to realize basic functions for face recognition. Therefore, in many situations, the difference between faces and objects is ‘quantitative’ rather than ‘qualitative’, as they are represented in a continuum of the representational space. In addition, we also found that face-specific experiences likely fine-tuned the face representation, and thus transformed the ‘object-like’ face processing into ‘face-specific’ processing. However, we shall be cautious that our finding may not be applicable for the development of face module in human, as in the biological brain experience-induced changes are partly attributed to the inhibition from lateral connections (Grill-Spector et al., 2006; Norman & O’Reilly, 2003), whereas there is no lateral or feedback connection in DCNNs. However, despite structural differences, recent studies have shown similar representation for faces between DCNNs and humans (Song, Qu, Xu, & Liu, 2020), suggesting that a common mechanism may be shared by both artificial and biological intelligent systems. Future studies are needed to examine the applicability of our finding to humans. On the other hand, our study illustrated the advantages of using DCNN as a model to understand human mind because of its computational transparency and its dissociation of factors in nature and nurture. Thus, our study invites future studies with DCNNs to understand the development of domain specificity in particular and a broad range of cognitive modules in general.

## 4 Materials and Methods

### 4.1 Stimuli

#### Deprivation dataset

The deprivation dataset was constructed to train the d-AlexNet. It was based on the ImageNet Large-Scale Visual Recognition Challenge (ILSVRC) 2012 dataset (Deng et al., 2009), which contains 1,281,167 images for training and 50,000 images for validation, in 1000 categories. These images were first subjected to automated screening with an in-house face-detection toolbox based on VGG-Face (Parkhi, Vedaldi, & Zisserman, 2015), and then further screened by two human raters, who separately judged whether a given image contains faces of humans or non-human primates regardless of the orientation and intactness of the face, or anthropopathic artwork, cartoons, and artifacts. We removed images judged by either rater as containing any above-mentioned contents. Finally, we removed categories whose remaining images were less than 640 images (approximately half of the original number of images in a category). The resultant dataset consists of 736 categories, with 662,619 images for training and 33,897 for testing the performance.

#### Classification dataset

To train a classifier that can classify faces, we constructed a classification dataset consisting of 204 categories of non-face objects and one face category, each of 80 exemplars. For the non-face categories, we manually screened Caltech-256 (Griffin, Holub, & Perona, 2007) to remove images containing human, primate, or cartoon faces, and then removed categories whose remaining images were less than 80. In each of the 204 remaining non-face categories, we randomly chose 70 images for training and another 10 for calculating classification accuracy. The face category was constructed by randomly selecting 1000 faces images from Faces in the Wild (FITW) dataset (Berg, Berg, Edwards, & Forsyth, 2005). Among them, 70 were used as training data and another 10 for classification accuracy. In addition, to characterize DCNN’s ability in differentiating faces from object categories, we compiled a second dataset consisting of all images in the face category except those used in training.

#### Discrimination dataset

To train a classifier that can discriminate faces at individual level, we constructed a discrimination dataset consisting of face images of 133 individuals, 300 images each, selected from the Casia-WebFace database (Yi, Lei, Liao, & Li, 2014). For each individual in the dataset, 250 were randomly chosen for training and another 50 for calculating discrimination accuracy.

#### Representation dataset

To examine representational similarity of faces and non-face images between the d-AlexNet and the normal one, we constructed a representation dataset with two categories, faces and bowling pins as an ‘unseen’ non-face object category that was not presented to the DCNNs during training. Each category consisted of 80 images. The face images were a random subset of FITW, and images of bowling pins were randomly chosen from the corresponding category in Caltech-256.

#### Movies clips for DCNN-brain correspondence analysis

We examined the correspondence between the face-selective response of the DCNNs and brain activity using a set of 18 clips of 8-min natural color videos from the Internet that are diverse yet representative of real-life visual experiences (Wen et al., 2017).

### 4.2 The deep convolutional neural network

Our model of selective deprivation, the d-AlexNet, was built with the architecture of the well-known DCNN ‘AlexNet’ (Krizhevsky et al., 2012, see Figure 1a for illustration). AlexNet is a feed-forward hierarchical convolutional neural network consisting of five convolutional layers (denoted as Conv1 – Conv5, respectively) and three fully connected layers denoted as FC1 – FC3. Each convolutional layer consists of a convolutional sublayer, followed by a ReLU sublayer, and Conv1, 2, and 5 are further followed by a pooling sublayer. Each convolutional sublayer consists of a set of distinct channels. Each channel convolves the input with a distinct linear filter (kernel) which extracts filtered outputs from all locations within the input with a particular stride size. FC1 to FC3 are fully connected layers. FC3 is followed by a sublayer using a softmax function to output a vector that represents the probability of the visual input containing the corresponding object category (Krizhevsky et al., 2012).

The d-AlexNet used the architecture of AlexNet but changed the number of units in FC3 to 736, so was the following softmax function, to match the number of categories in the deprivation dataset. Same to the pre-trained AlexNet in pytorch 1.2.0 (https://pytorch.org/, Paszke et al., 2017), the d-AlexNet was initialized with values drawn from a uniform distribution, and was then trained on the deprivation dataset following the approach specified in Krizhevsky et al., (2014). We used the pre-trained AlexNet from pytorch 1.2.0 as the normal DCNN, referred to as the AlexNet in this paper for brevity.

The present study referred to channels in the convolutional sublayers by the layer they belong to and a channel index, following the convention of pytorch 1.2.0. For instance, Layer 5-Ch256 refers to the 256^th^ convolutional channel of Layer 5.

### 4.3 Transfer learning for classification and discrimination

To examine to what extent our manipulation of the visual experience affected the categorical processing of faces, we replaced the fully-connected layers of each DCNN with a two-layer face-classification classifier. The first layer was a fully connected layer with 43,264 units as inputs and 4,096 units as outputs with sigmoid activation function, and the second was a fully connected layer with 4,096 units as inputs and 205 units as outputs, each of which corresponded to one category of the classification dataset. This classifier, therefore, classified each image into one category of the classification dataset. The face-classification classifier was trained for each DCNN with the training images in the classification dataset for 90 epochs.

To examine to what extent our manipulation of the visual experience affected face discrimination, we similarly replaced the fully connected layers of each DCNN with a discrimination classifier. The discrimination classifier differed from the classification classifier only in its second layer, which had 133 units instead as outputs, each corresponding to one individual in the discrimination dataset. The face-discrimination classifier was trained for each DCNN with the training images in the discrimination dataset for 90 epochs.

### 4.4 The face selective channels in DCNNs

To identify the channels selectively responsive to faces, we submitted images in the classification dataset to each DCNN, recorded the average activation in each channel of Conv5 after ReLU in response to each image, and then averaged the channel-wise activation within each category. We selected channels where the face category evoked the highest activation, and used the Mann-Whitney U test to examine the activation difference between faces and objects that had the second-highest activation in these channels (*p* < .05, Bonferroni corrected). The selectivity of each face channel thus identified was indexed by the selective ratio. The selective ratio was calculated by dividing the face activation by the second-highest activation. In addition, we measured the lifetime sparseness of each face-selective channel as an index for selectivity of faces among all non-face objects. We first normalized the mean activations of a face channel in Layer5 to all the categories to the range of 0-1, and then calculated lifetime sparseness with the formula:

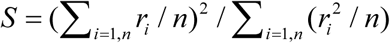

where r_i_ is the normalized activations to the ith object category. The smaller this value is, the higher the selectivity is.

### 4.5 DCNN-Brain Correspondence

We submitted the movie clips to the DCNNs. Following Wen (2017)’s approach, we extracted and log-transformed the channel-wise output (the average activation after ReLU) of each face-selective channel using the toolbox DNNBrain (Chen et al., 2020), and then convolved it with a canonical hemodynamic response function (HRF) with a positive peak at 4s. The HRF convolved channel-wise activity was then down-sampled to match the sampling rate of functional magnetic resonance imaging (fMRI) and the resultant timeseries was standardized before further analysis.

Neural activation in the brain was derived from the preprocessed data in Wen (2017). The fMRI data were recorded while human participants viewed each movie clips twice. We averaged the standardized time series across repetition and across subjects for each clip. Then, for each DCNN, we conducted multiple regression for each clip, with the activation time series of each brain vertex as the dependent variable and that of face-selective channels in this network as independent variables. For the d-AlexNet, all face-selective channels were included. For the AlexNet, we included the same number of face-selective channels with the highest face selectivity to match the complexity of the regression model. We used the R^2^ of each vertex as the index of the overall Goodness of fit of the regression in that vertex. The R^2^ values were then averaged across clips. The larger the R^2^ value, the higher correspondence between the DCNN and the brain in response to movie clips.

To determine whether cortical regions with large R^2^ values were traditional face-selective regions, we delineated the bilateral fusiform face areas (FFA) and the occipital face area (OFA) with the maximum-probability atlas of face-selective regions (Zhen et al., 2015). Two hundred of vertexes of the highest probability of the left FFA and 200 of the right FFA were included in the ROI of FFA, and the ROI of OFA was delineated in the same way. The correspondence with brain activation in each ROI and the impact of the visual experience was examined by submitting the vertex-wise R^2^ into a two-way ANOVA with visual experience (d-AlexNet vs. AlexNet) as within-subject factor and regional correspondence (OFA and FFA) as between-subject factor.

### 4.6 Face inversion effect in DCNNs

The average activation amplitude of the top 2 face-selective channels of each DCNN in response to upright and inverted version of 20 faces from the Reconstructing Faces dataset (VanRullen & Reddy, 2019) was measured. The inverted faces were generated by vertically flipping the upright ones. The face inversion effect in the d-AlexNet was measured with paired sample t-tests (two-tailed) and the impact of the experience on the face inversion effect was examined by two-way ANOVAs with visual experience (d-AlexNet vs. AlexNet) and inversion (upright vs inverted) as within-subject factors.

### 4.7 Representational similarity analysis

To examine whether faces in the d-AlexNet were processed in an object-like fashion, we compared the within-category representational similarity of faces to that of bowling pins, an ‘unseen’ non-face object category never exposed to either DCNN. Specifically, for each image in the representation dataset, we arranged the average activations of each channel of Conv5 after ReLU into vectors, and then for each pair of images we calculated and then Fisher-z transformed the correlation between their vectors, which served as an index of pairwise representational similarity. Within-category similarity between pairs of face images and that between pairs of object images were calculated separately. A 2 × 2 ANOVA was conducted with visual experience (d-AlexNet vs AlexNet) and category (face vs object) as independent factors. In addition, cross-category similarity between faces and bowling pins was also calculated for each DCNN, and a paired sample t-test (two-tailed) on two DCNNs was conducted.

### 4.8 Sparse coding and empirical receptive field

To quantify the degree of sparseness of the face-selective channels in representing faces, we submitted the same set of 20 natural images containing faces from FITW to each DCNN, and measured the number of activated units (i.e., the units showing above-zero activation) in the face-selective channels. The more non-zero units observed in the face-selective channels, the less sparse the representation for faces is. The coding sparseness of the two DCNNs was compared with a paired-sample t-test.

We also calculated the size of the empirical receptive field of the face-selective channels. Specifically, we obtained the activation maps of 1000 images randomly chosen from FITW. Using the toolbox DNNBrain (Chen et al., 2020), we up-sampled each activation maps to the same size of the input. For each image, we averaged the up-sampled activation within the theoretical receptive field of each unit (the part of the image covered by the convolution of this unit and the preceding computation, decided by the network architecture), and selected the unit with the highest average activation. We then cropped the up-sampled activation map by the theoretical receptive field of this unit, to locate the image part that activated this channel most across all the units. Then, we averaged corresponding cropped activation maps across all the face images, and the resultant map denotes the empirical receptive field of this channel, delineating the part of the theoretical receptive field that causes this channel to respond strongly in viewing its preferred stimuli.

## 5 Acknowledgement

This study was funded by the National Natural Science Foundation of China (31861143039, 31771251, and 31600925), the Fundamental Research Funds for the Central Universities, and the National Basic Research Program of China (2018YFC0810602).

## 6 Conflict of Interest

The authors declare no competing interests.

## Notes

### Competing Interest Statement

The authors have declared no competing interest.

